# SPARC: a method to genetically manipulate precise proportions of cells

**DOI:** 10.1101/788679

**Authors:** Jesse Isaacman-Beck, Kristine C. Paik, Carl F. R. Wienecke, Helen H. Yang, Yvette E. Fisher, Irving E. Wang, Itzel G. Ishida, Gaby Maimon, Rachel I. Wilson, Thomas R. Clandinin

**Affiliations:** Department of Neurobiology, Stanford University, Stanford, CA 94305; Department of Medicine, Weill Cornell Medical College, New York, NY 10065; Department of Neurobiology and Howard Hughes Medical Institute, Harvard Medical School, Boston, MA 02115; Freenome, South San Francisco, CA 94080; Laboratory of Integrative Brain Function and Howard Hughes Medical Institute, The Rockefeller University, New York, NY 10065

## Abstract

Many experimental approaches rely on controlling gene expression in select subsets of cells within an individual animal. However, reproducibly targeting transgene expression to specific fractions of a genetically-defined cell-type is challenging. We developed <u>S</u>parse <u>P</u>redictive <u>A</u>ctivity through <u>R</u>ecombinase <u>C</u>ompetition (SPARC), a generalizable toolkit that can express any effector in precise proportions of post-mitotic cells in *Drosophila*. Using this approach, we demonstrate targeted expression of many effectors and apply these tools to calcium imaging of individual neurons and optogenetic manipulation of sparse cell populations *in vivo*.

## Main Text

Genetic labeling and manipulation of small groups of cells in somatic mosaic animals have provided significant insights into many aspects of biology and have been particularly impactful in studies of the nervous system. Indeed, measurement and manipulation of genetically defined cell types has become central to neural circuit dissection in many systems^1^. Especially powerful are paradigms in which one measures the phenotypes of stochastically selected individual cells within otherwise unmanipulated populations. However, few methods exist for selectively manipulating a desired fraction of cells of the same genetically-defined type. In rodents, sequential recombinase-mediated switches can label subpopulations of neurons, but require labor-intensive titration of viruses^2^. In *Drosophila*, GAL4 and split GAL4 lines enable targeting of single cell types^3^, but selective manipulation of subsets of neurons of the same type remains challenging. For example, effector expression can be restricted by limiting the spatial and/or temporal expression of a recombinase, but this necessitates user-dependent heat shock or chemical induction, and in some cases, cannot be used in post-mitotic cells^4-6^. Therefore, a routine all-genetic method of expressing effectors in defined fractions of post-mitotic cells of the same type would provide a powerful means of dissecting cellular and genetic functions.

To address this need, we developed SPARC, a toolkit to express any effector in precise proportions of post-mitotic cells labelled by the GAL4-UAS system (Fig. 1A, S1, S2)^7^. The core of this toolkit is a set of bistable UAS-driven constructs that can be switched on or off in different relative proportions of cells, depending on their sequences. We designed each UAS-driven construct, so that PhiC31 recombinase^8^ could irreversibly recombine one of two competing *attP* target sequences with one *attB* target sequence. The reaction mediated by the first *attP* would remove a stop cassette to enable effector expression in cells expressing Gal4, while the reaction using the second *attP* would leave this stop intact and prevent expression (Fig. S1). Truncating canonical *attP* sequences diminishes the efficacy of recombination *in vitro*^9^. Based on this, we reasoned that truncating the first *attP* relative to the second would shift the equilibrium to favor retention of the stop cassette and result in sparser effector expression *in vivo*.

**Figure 1:**
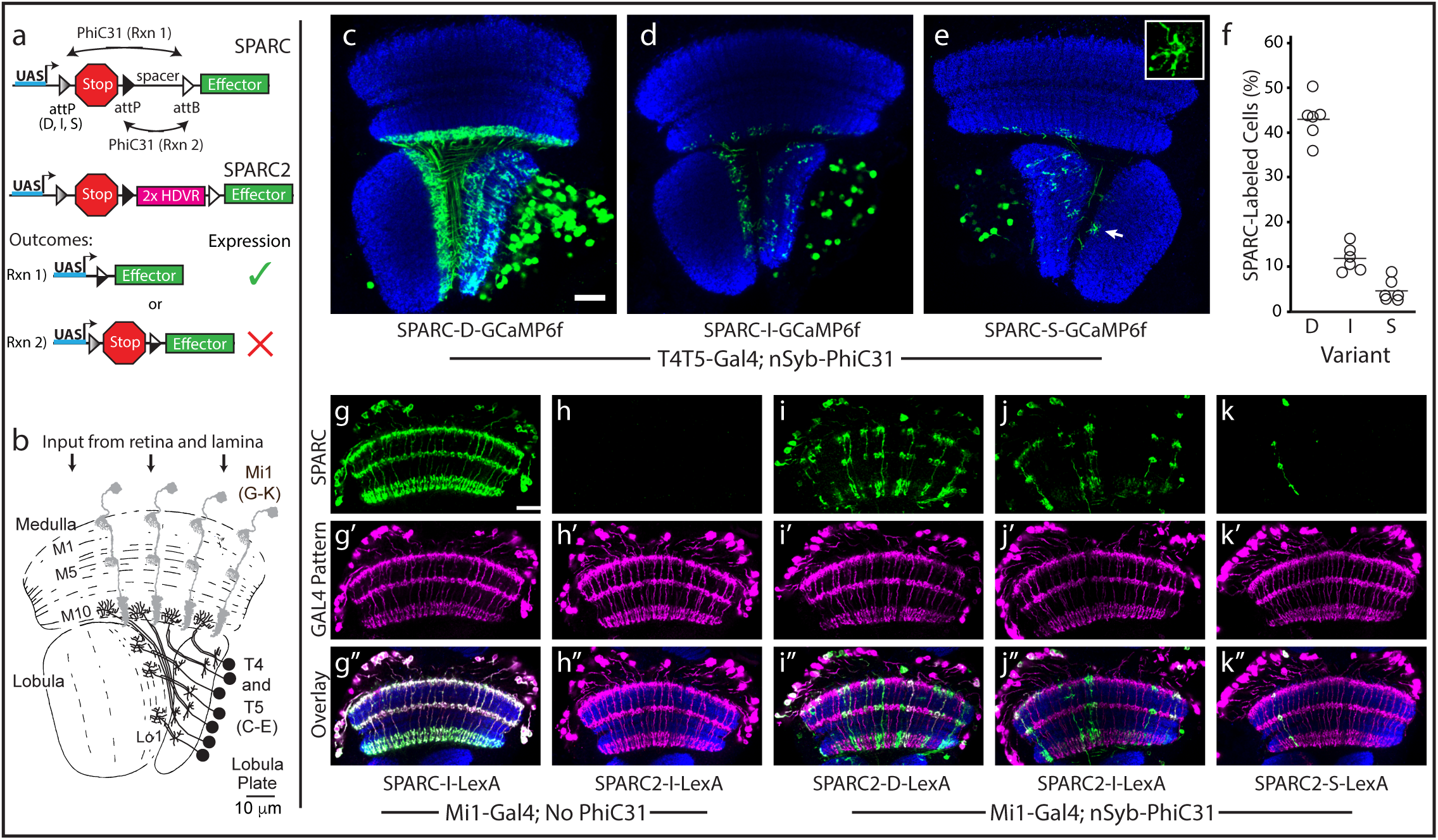
SPARC and SPARC2 enable predictive expression of effectors at three levels. **(a)** Schematic of SPARC and SPARC2 modules. PhiC31 recombines one of two competing attP target sequences with one attB target sequence. Truncating the first attP progressively favors retention of the stop cassette, preventing expression (Dense-60bp, canonical sequence; Intermediate - 38bp; Sparse – 34bp). SPARC2 incorporates a 2X Hepatitis Delta Virus Ribozyme (HDVR) sequence to prevent read-through in the absence of PhiC31. (b) Schematic of the Drosophila optic lobe highlighting T4, T5 and Mi1. (c-e) GCaMP6f expression (green) in T4 and T5 neurons counterstained with anti-Bruchpilot (blue). (c) SPARC-D-GCaMP6f (d) SPARC-I-GCaMP6f (e) SPARC-S-GCaMP6f, arrow points to dendrite shown in inset (f) Percentage of T4 and T5 neurons labeled by different SPARC modules (see Fig. S3). N = 6 optic lobes and > 400 cell bodies/genotype. (g-k) LexA::p65-driven expression of lexAop-myr-tDT (green, g-k), in Mi1 neurons (magenta, g’-k’) counterstained with anti-Bruchpilot (blue, overlay, g”-k”). (g) SPARC-I-LexA::p65 no PhiC31. LexA::p65 is expressed in all Mi1 neurons in the absence of PhiC31 in these animals. (h) SPARC2-I-LexA::p65 no PhiC31. LexA::p65 is not expressed in Mi1 neurons in the absence of PhiC31, consistent with an absence of read-through. (i) SPARC2-D-LexA::p65. (j) SPARC2-I-LexA::p65 (k) SPARC2-S-LexA::p65. Scale bars, 10 µm; N > 10 optic lobes per condition.

To test this concept, we generated plasmids and transgenic flies bearing SPARC constructs expressing the calcium indicator GCaMP6f^10^ and including one of three different *attP* variants at the first position (canonical: attP60; truncated: attp38 or attp34)^9^. We also generated transgenic flies that express PhiC31 recombinase in all post-mitotic neurons under the control of the synaptobrevin promoter^3^. We then drove expression of each SPARC construct in one of the largest genetically-defined populations of neurons in the *Drosophila* optic lobe, T4 and T5 cells^11^, in animals that express PhiC31 pan-neuronally. Across SPARC-GCaMP6f variants, we observed progressively fewer labeled neurons (Fig. 1C-E). SPARC-*attP60*-GCaMP6f labeled many overlapping neurons, SPARC-*attP38*-GCaMP6f labeled an intermediate number of neurons and SPARC-*attP34*-GCaMP6f labeled individual neurons whose dendrites could be visualized (Fig 1E inset). We therefore named these variants SPARC-D (Dense), SPARC-I (Intermediate), and SPARC-S (Sparse). To determine the percentage of these neurons labeled by each SPARC module, we expressed myristoylated-tdTomato (myr-tdT) in all T4 and T5 neurons in parallel with SPARC-GCaMP6f and counted singly and doubly labeled T4 and T5 cell bodies (Fig. 1F and S3). For SPARC-D-GCaMP6f we observed effector expression in ∼45% of T4 and T5 neurons. In comparison, SPARC-I-GCaMP6f labeled ∼12% of T4 and T5 neurons, while SPARC-S-GCaMP6f labeled only ∼4% of these neurons. We observed similar results in Kenyon cells, Lobula-Columnar neurons, and several columnar neurons in the optic lobe (Fig. S4 and data not shown). These data demonstrate that the SPARC module can reproducibly determine the fraction of cells that express effector over a more than 10-fold range across cells and animals. As SPARC-S can readily label T4, T5 and Kenyon cells, very common cell types in the fly brain, these studies argue that one of the three SPARC modules should allow targeting of individual cells of any cell type of interest.

To generalize this technique, we next made SPARC-LexA::p65 transgenes. LexA::p65 is a transcription factor that drives expression of transgenes under the control of the lexAop promoter^12^; this system is orthogonal to the UAS-GAL4 expression system. We expressed SPARC-LexA::p65 in Mi1 neurons of the *Drosophila* optic lobe (Fig 1B), and found that in the absence of PhiC31 recombinase, SPARC-LexA::p65 labels 100% of these neurons with lexAop-myr-tdT (Fig. 1G-G”). This result suggested that the widely-used stop cassette^13^ that we used in the initial SPARC design might permit a low level of read-through that can be detected by sensitive outputs like LexA::p65 (or mCD8::GFP, data not shown).

To avoid this read-through, we generated SPARC2, in which we incorporated two self-cleaving ribozymes from the Hepatitis Delta Virus (HDV) into the SPARC module (Fig. 1A). We reasoned that these self-cleaving ribozymes should truncate any read-through transcript prior to translation^14,15^. We first examined SPARC2-LexA::p65 transgenes in Mi1 neurons in the absence of PhiC31 and observed a 10,000-fold decrease in read-through (∼0.01% of Mi1 labeled with lexAop-myr-tdT, Fig. 1H-H”). Importantly, the D, I, and S variants of SPARC2-LexA::p65 behaved similarly to SPARC-GCaMP6f transgenes (Fig. 1I–K”). Also quantitative measurement of SPARC2-mCD8::GFP in Mi1 cells showed the same three levels of expression(Figure S4I-L). Thus, HDV ribozymes effectively eliminate read-through and enable SPARC2 transgenes to express both direct and amplifying effectors in three different proportions of cells.

Next, to investigate the functional utility of SPARC, we first used SPARC-S-GCaMP6f to image calcium (Ca^2+^) response in the dendrites of individual T5 neurons. These neurons preferentially respond to visual motion in one direction, a direction selectivity that is first observed in their dendrites^16^. Previous attempts to image from individual T5 cells relied on laborious FlpOut approaches that required titrated and temporally precise heat shocks of *Drosophila* larvae to restrict effector expression to a subset of cells^16-18^. In contrast, the SPARC method consistently labeled fewer T5 neurons, and labeled them more sparsely, than the sparsest FlpOut labeling using brief and developmentally late heat-shock (Fig. 1E, 2A,B). More importantly, when we imaged visually-evoked Ca^2+^ responses in regions of interest (ROIs) representing T5 dendrites, we observed that the fluorescent signals from SPARC-labeled ROIs were significantly more direction selective than those from FlpOut-labeled ROIs (DSI; Fig. 2C-E). This result reflects the fact that SPARC labeling was sparser than the sparsest FlpOut labeling we could achieve. As a consequence, SPARC ROIs more nearly captured signals from single cells, while FlpOut ROIs likely included signals from multiple labeled cells with different directional preferences (see supplemental methods). Thus, both anatomical and functional evidence suggests that SPARC better isolates single T5 dendrites more easily and more consistently than standard FlpOut approaches.

**Figure 2:**
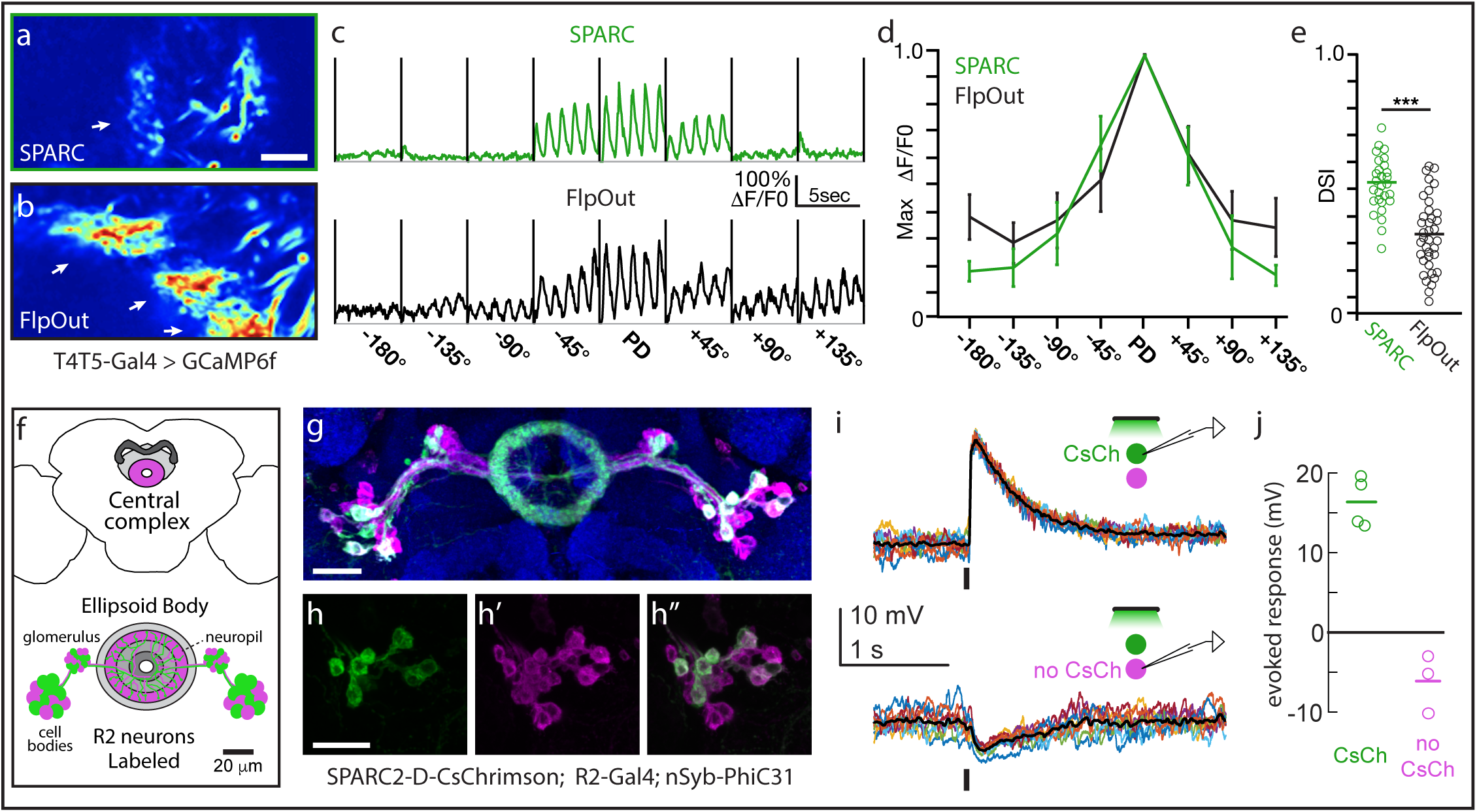
SPARC and SPARC2 enable calcium imaging of single neurons and optogenetic stimulation of sparse cell populations. **(a, b)** Normalized averaged fluorescence intensity of GCaMP6f in T5 dendrites sparsely labeled using (a) SPARC-S-GCaMP6f or (b) FlpOut-GCaMP6f. Arrows point to dendrites. **(c-d)** GCaMP6f fluorescence responses (ΔF/F0) of T5 dendritic ROIs to sinusoidal gratings moving in one of eight different directions. PD denotes the preferred direction of each cell with the angular deviation from PD in degrees (c) Averaged responses of a representative ROI labeled using SPARC-S-GCaMP6f (green) or FlpOut-GCaMP6f (black). (d) Normalized tuning curves averaged across all T5 dendritic ROIs labeled by SPARC-S-GCaMP6f (green) or FlpOut-GCaMP6f (black) **(e)** Direction selectivity indices (DSI) for each T5 dendritic ROI labeled by SPARC-S-GCaMP6f or FlpOut-GCaMP6f, n = 8 flies and 37 units per condition; *** p<0.001. **(f)** Schematic of Central Complex and Ellipsoid Body depicting SPARC2-labeled R2 Ring neurons (adapted from24). **(g)** SPARC2-D-CsChrimson::tdTomato-3.1 expression (tdTomato; green) in R2 Ring neurons (mCD8::GFP; magenta) counterstained with anti-Bruchpilot (blue). **(h-h”)** Closeup of cell bodies in (g) showing (h) CsChrimson expression in (h’) R2 cells. (h”) Overlay. **(I)** Current clamp recordings of single tdTomato+ (top) and tdTomato- (bottom) R2 neurons. Stimulus is a 50 ms pulse of green light; 10 trials each (colored lines), mean response (black line). **(j)** Average evoked response (open circles) and mean population response (line) of R2 neurons, both tdTomato+ (green, N =4) and tdTomato- (magenta, N = 3). Scale bars: 10 µm (a,b), 20 µm (g, h).

Then, to determine if we could use this approach to manipulate the activity of neuronal subpopulations, we generated SPARC2-CsChrimson::tdTomato^19^ transgenic flies. We expressed this construct in Ring (R) neurons, GABAergic neurons that send sensory input to the central complex^20^. R neurons are divided into types based on morphology^21^, here we focused on the R2 type. We expressed SPARC2-D-CsChrimson::tdTomato in a subset of R2 neurons (Figures 2F-H”). We observed that tdTomato^+^ R2 neurons were depolarized by light, while tdTomato^-^ R2 neurons were not depolarized (Figure 2I, J). Indeed, tdTomato^-^ R2 neurons were slightly hyperpolarized by light (Figure 2I, J), implying that these R2 neurons were postsynaptic to other R2 neurons that express CsChrimson. Thus, the SPARC system enables optogenetic activation of sparse cell populations within a single cell type, allowing us to discover evidence for mutual inhibitory interactions within a cell type. Although it was already known that some R neuron types inhibit other R types^20^, it was not previously known that there is mutual inhibition between R neurons of the same type.

The SPARC and SPARC2 toolkit includes direct effector transgenes that can be used to label cells (mCD8::GFP), to observe changes in intracellular calcium concentration (GCaMP6f, jGCaMP7f) and membrane potential (ASAP2f), as well as to optogenetically modulate neuronal activity (CsChrimson). In addition, the availability of the indirect effector transgene SPARC2-LexA::p65 opens the possibility of sparsely expressing a large range of additional existing effectors under the control of lexAop (Figure S5). To provide the flexibility to target both neuronal and non-neuronal cells, we also generated transgenic animals that express PhiC31 pan-neuronally (nSyb-PhiC31), ubiquitously (tub-PhiC31) and in any cell type labeled by GAL4 (UAS-PhiC31). Thus, by simply generating flies with the appropriate combination of transgenes (Fig. S6), one can perform a diverse array of experiments on single cells or precise proportions of cells of a given genetically defined type. Moreover, each element of this toolkit is modular, allowing users to easily incorporate any current or future genetically encoded effector (Fig. S3). In the context of the nervous system, SPARC, SPARC2 and future variations will allow convenient and unparalleled access to define the heterogeneity of single neuron contributions to neural circuit processing. In non-neuronal cells, SPARC will enable wide-ranging studies that exploit mosaic analysis to investigate the cell biology and physiology of post-mitotic cells. Finally, as PhiC31 functions in both the mouse and fish^22,23^, we anticipate that this strategy will be widely generalizable to other model systems.

## Acknowledgements

We thank members of the Clandinin, Wilson, and Maimon labs for discussion of the project and manuscript. We thank Arun Chakravorty for generating the SPARC-jGCaMP7f plasmids, Scott Gratz, Kate O’Connor-Giles, ChiChi Xie, Liqun Luo, Barrett Pfeiffer and David Anderson for providing template plasmids for molecular cloning. We also thank Gerald Rubin and Heather Dionne for sharing split-Gal4 stocks, Norbert Perrimon for sharing Cas9 stocks, and note that stocks obtained from the Bloomington Drosophila Stock Center (NIH P40OD018537) were used in this study. The project was supported by the NIH (R01EY022638 and 5P30EY026877 to T.R.C., 5U19NS104655 to T.R.C. and R.E.W. J.I-B. is an Arnold O. Beckman Postdoctoral Fellow, H.H.Y. is an HHMI fellow of the Jane Coffin Childs Memorial Fund for Medical Research, Y.E.F. is supported by a Hanna H. Grey Fellowship from HHMI, C.F.R.W. is supported by an NSF Graduate Research Fellowship (DGE – 1656518). R.I.W. and G.M. are HHMI investigators.

## Author contributions

J.I-B., H.H.Y, I.E.W. and T.R.C conceived the study. J.I-B, C.F.R.W., H.H.Y., and Y.E.F. designed and performed the experiments under the supervision of T.R.C. and R.I.W. J.I-B., K.C.P., H.H.Y., Y.E.F., and I.G.I generated, maintained, and/or validated transgenic fly stocks under the supervision of T.R.C, R.I.W., and G.M. J.I-B., C.F.R.W., H.H.Y., Y.E.F. and K.C.P. analyzed the data. J.I-B. and T.R.C. prepared the manuscript with contributions from C.F.R.W., H.H.Y., and Y.E.F.

## Methods

Attached as Supplemental Methods.

