## Supplemental Figures and Methods for "SPARC: a method to genetically manipulate precise proportions of cells"

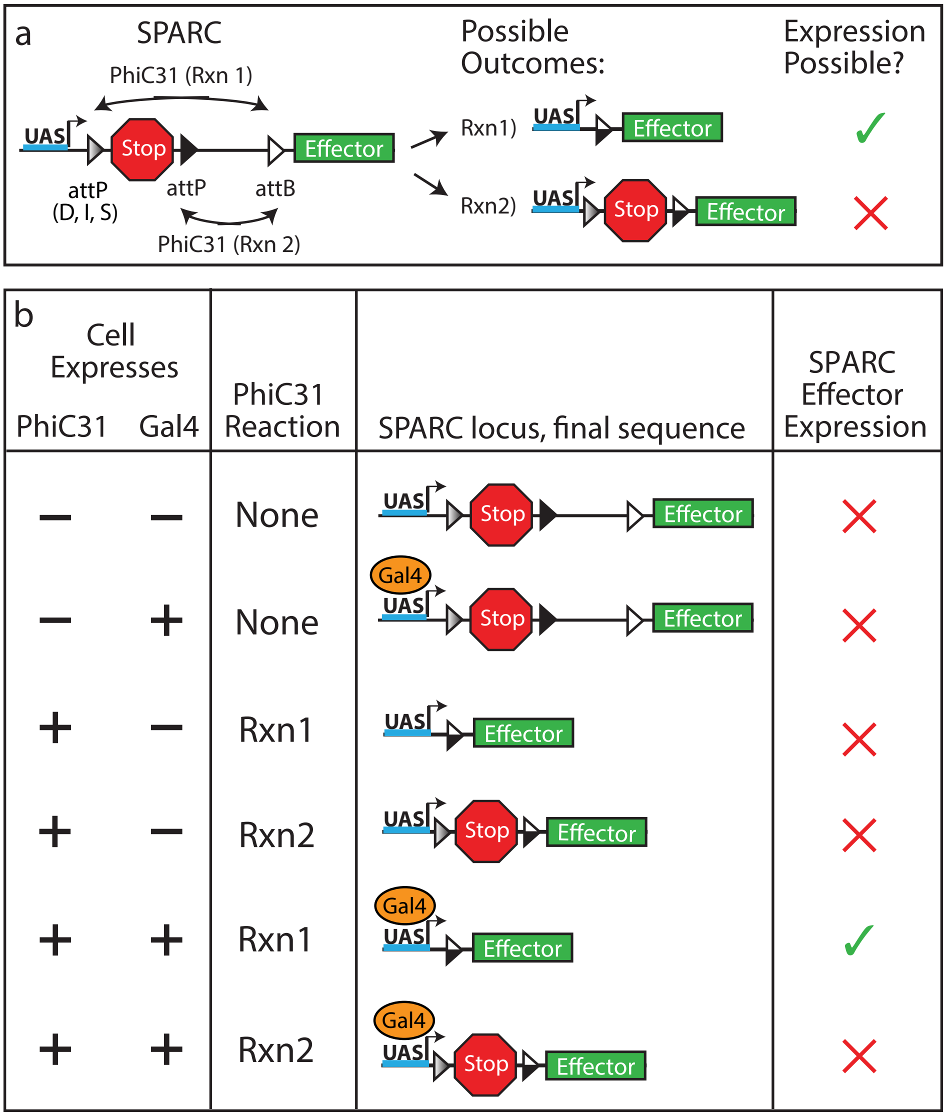


**Figure S1: Genetic conditions necessary for SPARC-Effector expression. (a)** PhiC31 irreversibly recombines SPARC module through one of two reactions that enables (Rxn 1) or prevents (Rxn 2) effector expression (Modified from Fig. 1A). **(b)** Table demonstrating how PhiC31 and GAL4 expression in a cell can impact the SPARC locus and SPARC-Effector expression. A cell must express both PhiC31 and GAL4 and PhiC31 must recombine the attP and attB sites for Reaction 1 to cause SPARC-effector expression.

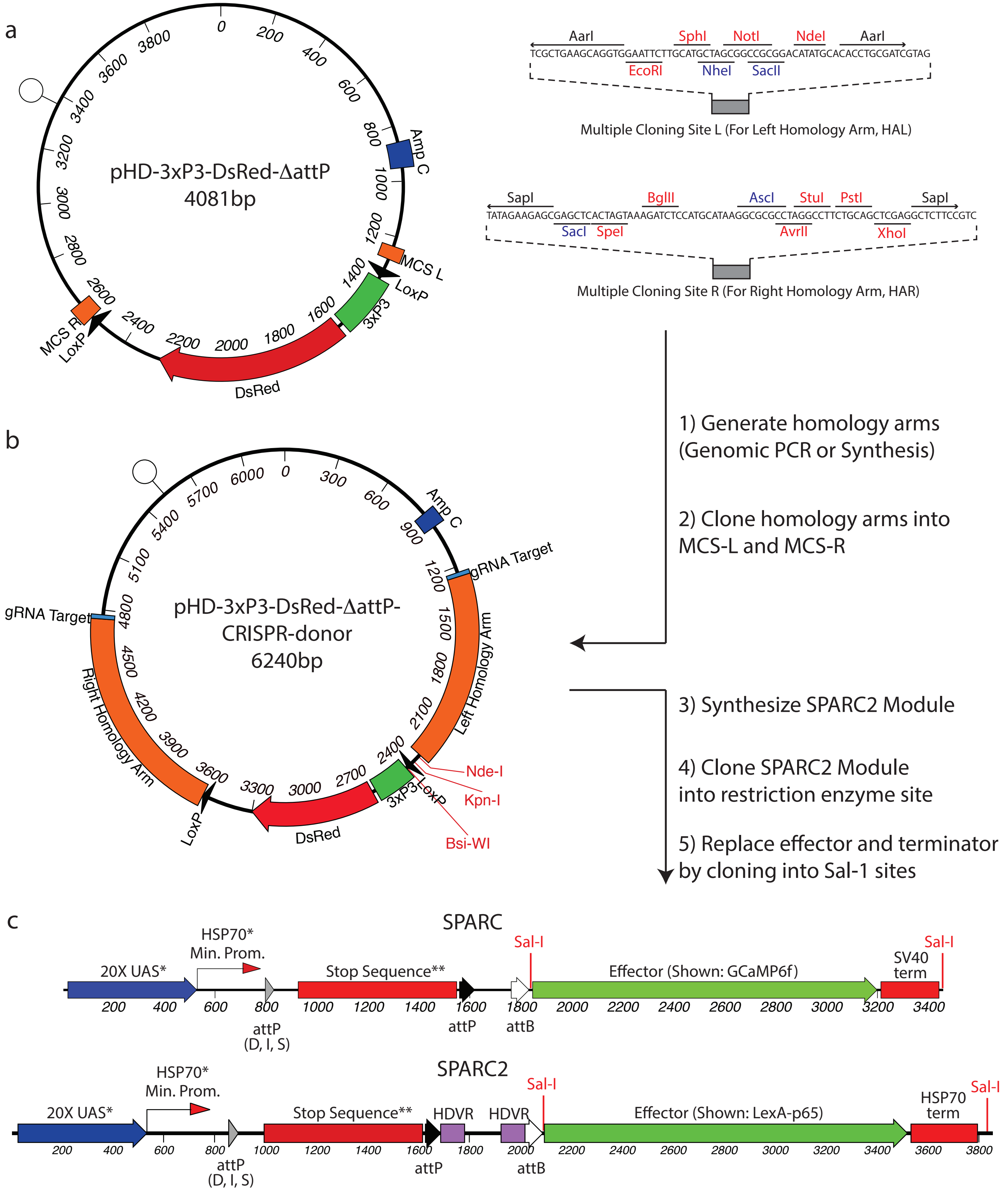


*Sequence for these elements was taken directly from pJFRC7 (Addgene #26220 ; Pfeiffer et al. (2010), Genetics, 186: 735-755.)

**Sequence for Stop Sequence was taken directly from pJFRC210 (Addgene #63170 ; Nern et al. (2015), Proc Natl. Acad. Sci., 112: E2967-E2976.)

Isaacman-Beck et al., 2019. Figure S2.

**Figure S2: Plasmid maps and molecular cloning methods for SPARC and SPARC2 constructs. (a)** Map of pHD-3xP3-DsRed-ΔattP (a CRISPR-HDR-donor precursor) showing multiple cloning sites for homology arm insertion (right). **(b)** Map of pHD-3xP3-DsRed-ΔattP-CRISPR-donor (example includes homology arms targeting attP40 region of the Drosophila genome). **(c)** SPARC and SPARC2 modules are inserted into pHD-3xP3-DsRed-ΔattP-CRISPR-donor via unique Kpn-I, Nde-I or Bsi-WI restriction enzyme sites. Sal-I restriction enzyme sites in the SPARC2 module allow for one-step swapping of the effector and terminator to generate pHD-SPARC2 donor plasmids. Abbreviations: MCS – multiple cloning site; gRNA – guide RNA; HDVR – Hepatitis Delta Virus Ribozyme sequence.

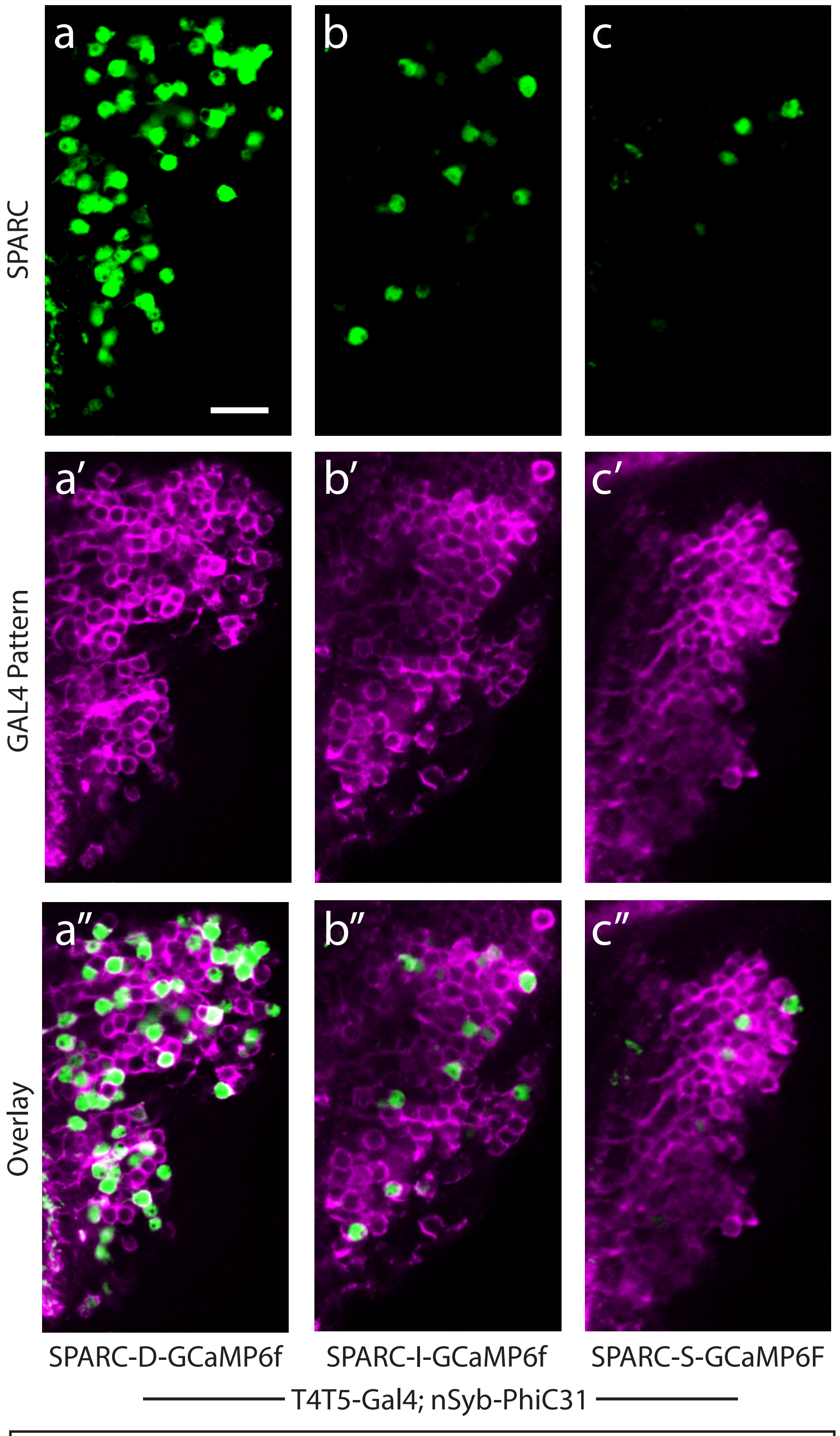


**Figure S3: Determining the proportion of T4 and T5 cells that express SPARC-GCaMP6f. (a-c″)** Single plane confocal images of GCaMP6f expression (green, a-c) in T4 and T5 cell bodies (magenta, a′-c′). Merge (a″-c″). (a-a″) SPARC-D-GCaMP6f. (b-b″) SPARC-I-GCaMP6f. (c-c″) SPARC-S-GCaMP6f.

Isaacman-Beck et al., 2019. Figure S3.

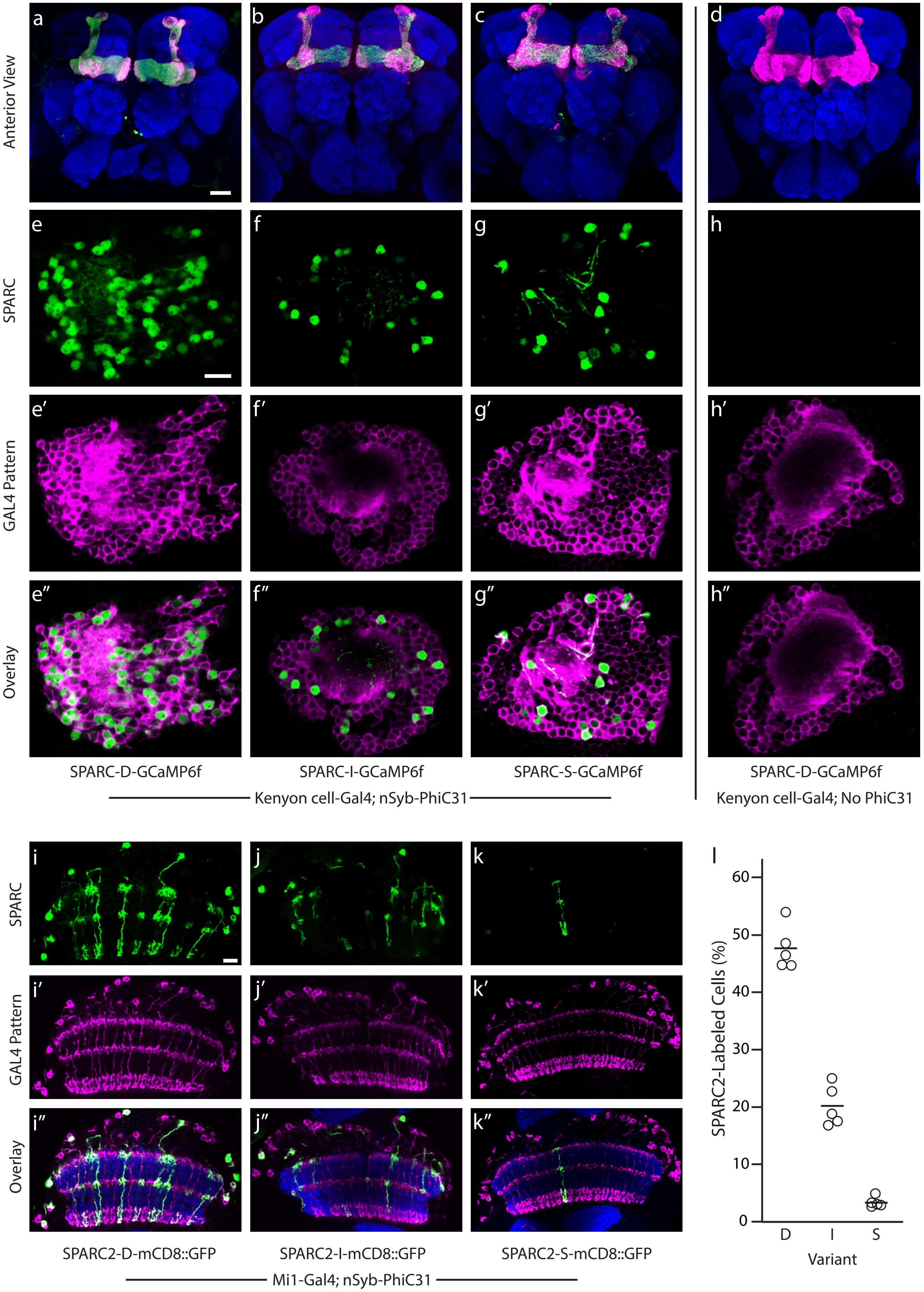


Isaacman-Beck et al., 2019. Figure S4.

**Figure S4: SPARC-GCaMP6f and SPARC2-mCD8::GFP expression in different cell types. (a-d)** Anterior view of *Drosophila* central brain showing GCaMP6f expression (green) in Kenyon cells (magenta) counterstained with anti-Bruchpilot (blue). (a) SPARC-D-GCaMP6f. (b) SPARC-I-GCaMP6f. (c) SPARC-S-GCaMP6f. (d) SPARC-D-GCaMP6f in the absence of PhiC31. **(e-h")** GCaMP6f expression (green, e-h) in Kenyon cell bodies (magenta, e'-h') with overlay (e"-h"). (e-e") SPARC-D-GCaMP6f. (f-f") SPARC-I-GCaMP6f. (g-g") SPARC-S-GCaMP6f. (h-h") SPARC-D-GCaMP6f in the absence of PhiC31. GCaMP6f is not detected in Kenyon Cells in the absence of PhiC31. **(i-k")** mCD8::GFP expression (green, i-k), in Mi1 neurons (magenta, i'-k') counterstained with anti-Bruchpilot (blue, overlay, i"-k"). (i-i") SPARC2-D-mCD8::GFP, (j-j") SPARC2-I-mCD8::GFP, and (k-k") SPARC2-S-mCD8::GFP. Scale bars = 30 µm (a-d), 10 µm (e-k"); N > 10 brains/condition (l) Percentage of Mi1 neurons labeled by different SPARC-mCD8::GFP modules. N = 5 optic lobes and > 300 cell bodies/genotype.

Transgenic Flies Generated in this Study

A. SPARC and SPARC2 Flies: Inserted with CRISPR-HDR into a region near the *attP40* locus (CR-P40; see supplemental methods). **Bolded stocks** will be available from the Bloomington Drosophila Stock Center (BDSC; bdsc.indiana.edu) ~November 1, 2019. Blue stocks will be available ~November 21, 2019 from BDSC.

1. **+; *TI{20XUAS-SPARC-D-GCaMP6f}CR-P40; +***
2. **+; *TI{20XUAS-SPARC-I-GCaMP6f}CR-P40; +***
3. **+; *TI{20XUAS-SPARC-S-GCaMP6f}CR-P40; +***
4. +; *TI{20XUAS-SPARC-D-jGCaMP7f}CR-P40; +*
5. +; *TI{20XUAS-SPARC-I-jGCaMP7f}CR-P40; +*
6. +; *TI{20XUAS-SPARC-S-jGCaMP7f}CR-P40; +*
7. **+; *TI{20XUAS-SPARC-I-ASAP2f}CR-P40; +***
8. +; *TI{20XUAS-SPARC-D-mCD8::GFP}CR-P40;* +
9. +; *TI{20XUAS-SPARC-I-mCD8::GFP}CR-P40;* +
10. +; *TI{20XUAS-SPARC-S-mCD8::GFP}CR-P40;* +
11. +; *TI{20XUAS-SPARC-D-LexA::p65}CR-P40;* +
12. +; *TI{20XUAS-SPARC-I-LexA::p65}CR-P40;* +
13. +; *TI{20XUAS-SPARC-S-LexA::p65}CR-P40;* +
14. **+; *TI{20XUAS-SPARC2-D-LexA::p65}CR-P40; +***
15. **+; *TI{20XUAS-SPARC2-I-LexA::p65}CR-P40; +***
16. **+; *TI{20XUAS-SPARC2-S-LexA::p65}CR-P40; +***
17. **+; *TI{20XUAS-SPARC2-D-Syn21-CsChrimson::tdTomato-3.1}CR-P40; +***
18. **+; *TI{20XUAS-SPARC2-I-Syn21-CsChrimson::tdTomato-3.1}CR-P40; +***
19. **+; *TI{20XUAS-SPARC2-S-Syn21-CsChrimson::tdTomato-3.1}CR-P40; +***
20. **+; *TI{20XUAS-SPARC2-D-mCD8::GFP}CR-P40; +***
21. **+; *TI{20XUAS-SPARC2-I-mCD8::GFP}CR-P40; +***
22. **+; *TI{20XUAS-SPARC2-S-mCD8::GFP}CR-P40; +***

B. PhiC31 transgenic strains:

1. ***y,w; P{nSyb-IVS-PhiC31}su(Hw)attP5; +***
2. ***y,w; P{20XUAS-IVS-PhiC31}su(Hw)attP5; +***
3. *y,w; P{αTub84-IVS-PhiC31}su(Hw)attP5; +*
4. ***y,w, P{nSyb-IVS-PhiC31}attP18; +***
5. ***y,w, P{20XUAS-IVS-PhiC31}attP18; +***
6. ***+, P{nSyb-IVS-PhiC31}attP18; s/cyo; pr/TM6B***
7. ***+, P{20XUAS-IVS-PhiC31}attP18; s/cyo; pr/TM6B***

**Figure S5: List of SPARC and SPARC2 transgenic flies generated in this study. (a)** SPARC and SPARC2 transgenic flies **(b)** PhiC31 transgenic flies generated in this study.

SPARC/SPARC2 User guide

A) Important notes:

1. SPARC and SPARC2 activity requires a minimum of 3 transgenes: 1) SPARC-effector, 2) promoter-PhiC31, and 3) enhancer-GAL4.
2. UAS-SPARC-Effector or UAS-SPARC2-Effector and promoter-PhiC31 transgenes must be maintained in seperate stocks to avoid germline recombination.
3. For all new UAS-SPARC-Effector combinations, we recommend using SPARC2.

B) Example crossing schemes for SPARC or SPARC2

$\frac{+}{+}$ ; SPARC2-Effector (near {attP40}) ; + X $\frac{+}{y}$ ; promoter*-PhiC31 {su(Hw)attP5} ; enhancer-GAL4 {attP2} OR

$\frac{+}{+}$ ; SPARC2-LexA::p65 (near {attP40}) ; lexAop-effector (variable) X $\frac{+}{y}$ ; promoter*-PhiC31 {su(Hw)attP5} ; enhancer-GAL4 {attP2}

* Available stocks: 20XUAS-IVS-PhiC31, atub84-IVS-PhiC31, and nSyb-IVS-PhiC31

Using Janelia split Gal4 Drivers:

$\frac{+}{+}$ ; SPARC2-effector (near {attP40} ; + X +, promoter*-PhiC31 {attp18} / y ; enhancer-GAL4AD {attP40} ; enhancer-GAL4DBD {attP2}

$\frac{+}{+}$ ; SPARC2-LexA::p65 (near {attP40} ; lexAop-effector (variable} X +, promoter*-PhiC31 {attp18} / y ; enhancer-GAL4AD {attP40} ; enhancer-GAL4DBD {attP2}

* Available stocks: 20XUAS-IVS-PhiC31 and nSyb-IVS-PhiC31

**Figure S6: SPARC and SPARC2 User Guide. (a)** Important notes regarding SPARC and SPARC2 use and stock maintenance. **(b)** Example crossing schemes for SPARC or SPARC2 to allow expression of effectors.

Isaacman-Beck et al., 2019. Figure S6.

### Supplemental Methods:

#### Generation of plasmids for transgenesis:

All plasmids were generated with In-Fusion cloning (Takara Biotech; Mountain View, CA) using the primers described below or were generated through synthesis and molecular cloning by Genscript (Piscataway, NJ, USA). Constructs were sequence-verified by single primer extension (Sequetech; Mountain View, CA). We have submitted the following constructs to Addgene:

| Plasmid Name | Function |
|---|---|
| pHD-3xP3-dsRed-ΔattP | Clone in homology arms for CRISPR-donor targeting |
| pHD-3XP3-dsRed-ΔattP-CRISPR-donor-attP40 | Clone in SPARC or SPARC2 modules for CRISPR-HDR genomic insertion near the attP40 locus |
| pCFD5-U6-3-t-attP40 | Guide RNA plasmid to co-inject with pHD-3XP3-dsRed-ΔattP-CRISPR-donor-attP40 for CRISPR-HDR |
| pHD-SPARC2-D-LexA::p65 | Swap out effector and terminator to generate SPARC2-D CRISPR-donor vector for genomic insertion near the attP40 locus |

| | |
|---|---|
| pHD-SPARC2-I-LexA::p65 | Swap out effector and terminator to generate SPARC2-I CRISPR-donor vector for genomic insertion near the attP40 locus |
| pHD-SPARC2-S-LexA::p65 | Swap out effector and terminator to generate SPARC2-S CRISPR-donor vector for genomic insertion near the attP40 locus |
| pJFRC-nSyb-IVS-PhiC31 | Replace the nSyb promoter to express PhiC31 under the control of any promoter of interest. |

All other constructs are available upon written request.

**PhiC31 recombinase construct synthesis:**

We generated three constructs to express PhiC31 recombinase under the control of different promoters: 20XUAS, Tubulin (Tub), and Synaptobrevin (nSyb). These constructs were built in the backbone of pJFRC7 Addgene #26220[13]. To generate pJFRC-20XUAS-IVS-PhiC31, we PCR-amplified a Kozak consensus sequence followed by the PhiC31 recombinase open reading frame and an NLS sequence from vector pBS130[25] and cloned it into the backbone via Xho-I and Xba-I sites. To generate pJFRC-$\alpha$Tub84B-IVS-PhiC31, we PCR-amplified the Tubulin 1 alpha ($\alpha$Tub84B) promoter from pIDT-attB-Tub_NLSGal4DBDlink-PBT[25] and replaced the 20XUAS promoter in pJFRC-20XUAS-IVS-PhiC31 by cloning into HindIII-BglII sites. To generate pJFRC-nSyb-IVS-PhiC31, the synaptobrevin (nSyb) promoter was PCR-amplified from the pattB-synaptobrevin-4-QFBDMD-G4AD-hsp70 plasmid[26] and cloned into the pJFRC-αTub84B -IVS-PhiC31 backbone via BglII sites flanking the αTub84B promoter. We submitted the pJFRC-nSyb-IVS-PhiC31 plasmid to Addgene (see table above).

**UAS-SPARC and UAS-SPARC2 CRISPR donor plasmid synthesis:**

To generate the SPARC and SPARC2 backbone plasmid, pHD-3xP3-dsRed-ΔattP, we used site-directed mutagenesis (Quikchange, Stratagene XL; Agilent, Santa Clara, CA, USA) to replace the attP sequence in the CRISPR donor vector pHD-AttP-3XP3-dsRed[27] with unique Kpn-I and Mlu-I restriction enzyme sites using primer pair 1 (below). We prepared this vector to target the attP40 region of the genome by PCR-amplifying a 1040 base pair (bp) left homology arm (2L:5106650..5107689) and an 1168bp right homology arm (2L:5108423..5109590) from genomic DNA of the Iso-D1 strain[28]. Left and right homology arms were first cloned into PCR2.1d-topo (Invitrogen, Waltham, MA, USA) using primer pairs 2 and 4 (below) and were then subcloned into pHD-3XP3-dsRed-ΔattP using primer pair 3 via Not-I and primer pair 5 via Sap-I respectively. Primer pair 3 and 5 added external flanking guide RNA (gRNA) target sites to enable Cas9-mediated linearization of the donor sequence *in vivo* (see Fig. S2).

**Primer pairs for CRISPR donor plasmid synthesis:**

| Primer Name | Pair | Sequence |
|---|---|---|
| Q5SDM_pHD_F4 | 1 | ATCGCGGTACCGGGGGCGTAGATAACTTC |
| Q5SDM_pHD_R1 | 1 | CGACTACGCGTCGATCGCAGGTGTGCATATG |
| HAL_attP40_F1 | 2 | TGTTTGCCAAAAGTGCTGGAG |

| | | |
|---|---|---|
| HAL_attP40_R1 | 2 | GCTGGTGGTTGTTTATGCCT |
| HAL_attP40_Not1_gRNA_F1 | 3 | TGCATGCTAGCGGCCGCACAGAAAAGCATCTTACGGATGGTGTTTGCCAAAAGTGCTGGAG |
| HAL_attP40_Not1_ R1 | 3 | CATATGTCCGCGGCCGCGCTGGTGGTTGTTTATGCCT |
| HAR_attP40_F1 | 4 | CACATTGGGTAGCACAGACTC |
| HAR_attP40_R1 | 4 | CGTACTCATCCTGGTCGTCG |
| HAR_attP40_SapI_F1 | 5 | TTCTGACCTGGGTATCACATTGGGTAGCACAGACTC |
| HAR_attP40_SapI_gRNA_R1 | 5 | CTTTCCGATATCGACACAGAAAAGCATCTTACGGATGG CGTACTCATCCTGGTCGTCG |

Next, several SPARC and SPARC2 modules were synthesized by Genscript (Piscataway, NJ, USA) and cloned into the unique Kpn-I site (Fig. S2).

**First UAS-SPARC and UAS-SPARC2 module plasmids:**

| SPARC Module | First attP sequence* | Effector |
|---|---|---|
| SPARC-D-GCaMP6f | attP60 | GCaMP6f[10] |
| SPARC-I-GCaMP6f | attP38 | GCaMP6f |
| SPARC-S-GCaMP6f | attP34 | GCaMP6f |
| SPARC-D-LexA::p65 | attP60 | LexA::p65[12] |
| SPARC-I-LexA::p65 | attP38 | LexA::p65 |

| | | |
|---|---|---|
| SPARC2-D-LexA::p65 | attP60 | LexA::p65 |
| SPARC2-I-LexA::p65 | attP38 | LexA::p65 |
| SPARC2-S-LexA::p65 | attP34 | LexA::p65 |
| SPARC-D-GCaMP7f | attP60 | jGCaMP7f[29] |
| SPARC-I-GCaMP7f | attP38 | jGCaMP7f |

* Sequences taken from[8].

Subsequently, the following donor constructs were generated by PCR cloning of new effectors into the appropriate SPARC or SPARC2 module backbone via In-Fusion cloning with Sal-I restriction digest (Clontech, Mountain View CA) or using the CloneEZ method (Genscript, Piscataway NJ). Constructs generated in this manner include:

**Second UAS-SPARC Module plasmids:**

| SPARC Module | First attP sequence | Effector |
|---|---|---|
| SPARC-S-LexA::p65 | attP34 | LexA::p65 |
| SPARC-D-mCD8::GFP | attP60 | mCD8::GFP[4] |
| SPARC-I-mCD8::GFP | attP38 | mCD8::GFP |
| SPARC-S-mCD8::GFP | attP34 | mCD8::GFP |
| SPARC-I-ASAP2f | attP38 | ASAP2f[30] |
| SPARC2-D-mCD8::GFP | attP60 | mCD8::GFP |
| SPARC2-I-mCD8::GFP | attP38 | mCD8::GFP |
| SPARC2-S-mCD8::GFP | attP34 | mCD8::GFP |

| | | |
|---|---|---|
| SPARC2-D-Syn21-CsChrimson::tdTomato-3.1 | attP60 | CsChrimson::tdTomato[31,32] |
| SPARC2-I-Syn21-CsChrimson::tdTomato-3.1 | attP38 | CsChrimson::tdTomato |
| SPARC2-S-Syn21-CsChrimson::tdTomato-3.1 | attP34 | CsChrimson::tdTomato |

For detailed construct maps of SPARC and SPARC2 as well as recommended methods for modular swapping of effector, see Fig. S2.

**gRNA-targeting vector logic and synthesis:**

Name of gRNA targeting vector: pCFD5-U6-3-t-attP40

We defined gRNA targets for insertion around the attP40 region of the genome using the publicly available search tool: flybase.org/crispr/[33]. The sequence and genomic location of these target sites as well as the synthetic gRNA used for donor plasmid linearization are as follows:

| gRNA Target | Genomic Location | Sequence | Strand |
|---|---|---|---|
| attP40_1 | 2L:5107682..5107704 | CCACCAGCCAAGGGCTGTGTAAC | Minus |
| attP40_2 | 2L:5108417..5108439 | CCAACACACATTGGGTAGCACAG | Minus |

| | | | |
|---|---|---|---|
| Synthetic gRNA | Not applicable | CCATCCGTAAGATGCTTTTCTGT | Minus |

We validated that these gRNA targets were present and unmutated in our CRISPR target flies (y,sc,v; + ; Nos-Cas9 (attP2), TH00787; kind gift from Norbert Perrimon) by PCR amplification and sequencing. Finally, we generated the construct pCFD5-attP40-gRNA using previously described methods (http://www.crisprflydesign.org/plasmids/) by PCR-amplifying three gRNA components and inserting them into the pCFD5: U6:3-t::gRNA backbone (a gift from Simon Bullock, Addgene #73914) [34] via Bbs-I (primers below).

| Name | Sequence |
|---|---|
| attP40_PCR_F1 | TTCCCGGCCGATGCAGTTACACAGCCCTTGGCTGGGTTTTAGAGCTAGAAATAGCAAG |
| attP40_PCR_R1 | ACACACATTGGGTAGCACAGTGCACCAGCCGGGAATCGAACCC |
| attP40_PCR_F2 | CTGTGCTACCCAATGTGTGTGTTTTAGAGCTAGAAATAGCAAG |
| attP40_PCR_R2 | TTCTAGCTCTAAAACTCCGTAAGATGCTTTTCTGTTGCACCAGCCGGGAATCGAACCC |

Two of these gRNAs targeted neighboring genomic regions near the attP40 genomic locus in our CRISPR-HDR target flies. We added a third gRNA component that targeted the synthetic gRNA sequence that flanks the donor insertion sequence in our pHD donor vectors (Fig. S2). This synthetic gRNA has no predicted off-target recognition in the *Drosophila melanogaster* genome[33].

**Generation of transgenic flies:**

All Phi31-expressing transgenic flies were generated using site-specific insertion into the genome[35]. These transgenes carry the miniW+ marker[36]; their genomic locations are listed in Fig. S5.

SPARC transgenic flies were generated by Bestgene (Chino Hills, CA, USA) via standard construct injections (100-500ng Donor construct, 75-250ng gRNA construct) and CRISPR-HDR. Transformants were identified by the marker 3xP3-DsRed which was later excised from the genome using Cre-Recombinase as previously described[27]. We maintained ≥3 independent transgenic flies for each transgene and tested them for expression and function.

**Genomic insertion site validation**:

To validate the insertion SPARC transgenes near the attP40 locus, we PCR amplified genomic DNA with primer pairs in which one primer recognizes genomic DNA beyond the homology arm and the other recognizes sequence within the transgene. For every SPARC or SPARC2 transgenic, we ensured amplification of PCR product from both the 5' and 3' sides of the transgene. The primer pairs are as follows:

| Primer Name | Sequence | Product Size (bp) |
|---|---|---|
| attP40-HAR-Verify-F1 | CTGGACATCACCTCCCACAAC | |
| attP40-HAR-Verify-R1 | ACATCGATCTCGAATGGATTCTCGG | ~1500bp |

| | | |
|---|---|---|
| attP40-HAL-Verify-F3 | GAGCGGAAAGCAATGTTTATGCGA | |
| attP40-HAL-Verify-R2 | CGGCCGCGCTGGTGGTTGTTT | ~1100bp |

**Note 1:** The attP40-HAR-Verify-F1primer recognizes sequence in the floxed 3xP3-DsRed sequence (see Fig. S2). Therefore, one must validate the insertion site using DNA from SPARC transgenic flies that still retain the floxed 3xP3-DsRed sequence.

**Note 2:** As HAL primers have some background amplification in control flies, we used a higher annealing temperature (~62ºC) for these amplifications.

All transgenic insertions were validated by PCR bands of the lengths noted above, with the exception of *TI{20XUAS-SPARC-D-Chrimson::tdTomato-3.1}CR-P40*. This transgene is missing ~300bp of the left homology arm. However, we validated the function of this transgene in Fig. 2.

**Complete Stock List:**

A list of all transgenic flies generated in this study can be found in Fig. S5.

**Transgenic genotypes (by Figure):**

Fig. 1C-E:

1. *+; TI{20XUAS-SPARC-D-GCaMP6f}CR-P40 / P{nSyb-IVS-PhiC31}su(Hw)attP5; P{VT025965-Gal4}attP2 / P{10XUAS-myr-tdTomato}attP2*
2. *+; TI{20XUAS-SPARC-I-GCaMP6f}CR-P40 / P{nSyb-IVS-PhiC31}su(Hw)attP5; P{VT025965-Gal4}attP2 / P{10XUAS-myr-tdTomato}attP2*

3. *+; TI{20XUAS-SPARC-S-GCaMP6f}CR-P40 / P{nSyb-IVS-PhiC31}su(Hw)attP5; P{VT025965-Gal4}attP2 / P{10XUAS-myr-tdTomato}attP2*

Fig. 1G-K:

1. *+; TI{20XUAS-SPARC-I-LexA::p65}CR-P40 /+; P{GMR19F01-Gal4}attP2 / P{UAS-mCD8::GFP}attP1, P{13XlexAop-IVS-myr-tdTomato}attP2*
2. *+; TI{20XUAS-SPARC2 -I-LexA::p65}CR-P40 /+; P{GMR19F01-Gal4}attP2 / P{10XUAS-mCD8::GFP}attP1, P{13XlexAop-IVS-myr-tdTomato}attP2*
3. *+; TI{20XUAS-SPARC2-D-LexA::p65}CR-P40 / P{nSyb-IVS-PhiC31}su(Hw)attP5; P{GMR19F01-Gal4}attP2 / P{10XUAS-IVS-mCD8::GFP}attP1, P{13XlexAop -myr-tdTomato}attP2*
4. *+; TI{20XUAS-SPARC2-I-LexA::p65}CR-P40 / P{nSyb-IVS-PhiC31}su(Hw)attP5; P{GMR19F01-Gal4}attP2 / P{10XUAS-IVS-mCD8::GFP}attP1, P{13XlexAop -myr-tdTomato}attP2*
5. *+; TI{20XUAS-SPARC2-S -LexA::p65}CR-P40 /P{nSyb-IVS-PhiC31}su(Hw)attP5; P{GMR19F01-Gal4}attP2 / P{10XUAS- IVS-mCD8::GFP}attP1, P{13XlexAop-myr-tdTomato}attP2*

Fig. 2A-E

1. *+; TI{20XUAS-SPARC-S-GCaMP6f}CR-P40 /P{nSyb-IVS-PhiC31}su(Hw)attP5; P{VT025965-Gal4}attP2/ +*
2. *+/y,w, P{hsFLP}; P{20XUAS-IVS-GCaMP6f}attP40/ P{αTub84b{FRT.Gal80}}2; P{VT025965-Gal4}attP2 /+*

Fig. 2G-J:

1. P{nSyb-IVS-PhiC31}attP18/+; TI{20XUAS-SPARC2-D-Syn21-CsChrimson::tdTomato-3.1}CR-P40 / {20XUAS-IVS-mCD8::GFP}attP40; P{GMR19C08-Gal4}attP2 / +
2. P{nSyb-IVS-PhiC31}attP18/y w; TI{20XUAS-SPARC2-D-Syn21-CsChrimson::tdTomato-3.1}CR-P40, 3XP3-DsRed/ P{20XUAS-IVS-mCD8::GFP}attP40; P{GMR19C08-Gal4}attP2 / +

Fig. S3:

4. +; TI{20XUAS-SPARC-D-GCaMP6f}CR-P40 /P{nSyb-IVS-PhiC31}su(Hw)attP5; P{VT025965-Gal4}attP2 / P{10XUAS-myr-tdTomato}attP2
5. +; TI{20XUAS-SPARC-I-GCaMP6f}CR-P40 /P{nSyb-IVS-PhiC31}su(Hw)attP5; P{VT025965-Gal4}attP2 / P{10XUAS-myr-tdTomato}attP2
6. +; TI{20XUAS-SPARC-S-GCaMP6f}CR-P40 /P{nSyb-IVS-PhiC31}su(Hw)attP5; P{VT025965-Gal4}attP2 / P{10XUAS-myr-tdTomato}attP2

Fig. S4A-H:

1. +; TI{20XUAS-SPARC-D-GCaMP6f}CR-P40 / P{nSyb-PhiC31}attP5, P{GMR19B03-Gal4AD}attP40 ; P{GMR026E07-Gal4DBD}attP2 / P{UAS-myr-tdTomato}attP2
2. +; TI{20XUAS-SPARC-I-SPARC-GCaMP6f}CR-P40 / P{nSyb-PhiC31}attP5, P{GMR19B03-Gal4AD}attP40 ; P{GMR026E07-Gal4DBD}attP2 / P{UAS-myr-tdTomato}attP2
3. +; TI{20XUAS-SPARC-S-GCaMP6f}CR-P40 / P{nSyb-PhiC31}attP5, P{GMR19B03-Gal4AD}attP40 ; P{GMR026E07-Gal4DBD}attP2 / P{UAS-myr-tdTomato}attP2

4. *+/w; TI{20XUAS-SPARC-D-GCaMP6f}CR-P40 / P{GMR19B03-Gal4AD}attP40 ; P{GMR026E07-Gal4DBD}attP2 / P{UAS-myr-tdTomato}attP2*

Fig. S4I-L:

1. *+; TI{20XUAS-SPARC2-D-mCD8::GFP}CR-P40 / P{nSyb-IVS-PhiC31}su(Hw)attP5; P{GMR19F01-Gal4}attP2 / P{UAS-myr-tdTomato}attP2*
2. *+; TI{20XUAS-SPARC2-I-mCD8::GFP}CR-P40 / P{nSyb-IVS-PhiC31}su(Hw)attP5; P{GMR19F01-Gal4}attP2 / P{UAS-myr-tdTomato}attP2*
3. *+; TI{20XUAS-SPARC2-S-mCD8::GFP}CR-P40 / P{nSyb-IVS-PhiC31}su(Hw)attP5; P{GMR19F01-Gal4}attP2 / P{UAS-myr-tdTomato}attP2*

**Origin of transgenes**

1. *P{GMR19F01-Gal4}attP2* from BDSC, stock number 48852, and is described in [3]
2. *P{VT025965-Gal4}attP2* from VDRC[37].
3. *P{GMR19B03-Gal4-AD}attP40; P{GMR026E07-Gal4DBD}attP2* from Janelia Flylight collection.
4. *P{GMR19C08-Gal4}attP2* from BDSC, stock number 48845, and is described in[3].
5. *P{10XUAS- myr-TdTomato}attP2* from BDSC, stock number 32221.
6. *P{13XlexAop- myr-TdTomato}attP2* was a generous gift from Heather Dionne and Gerald Rubin.
7. *P{10XUAS-IVS-mCD8::GFP}attP1* from BDSC, stock number 32187.
8. *P{20XUAS-IVS-mCD8::GFP}attP40* was a gift from Barret Pfeiffer and Gerald Rubin and is described in[13].

9. *CyO, P{w[+mC]=Crew}DH1* from BDSC, stock number 1092, and is described in[38].
10. *P{20XUAS-IVS-GCaMP6f}attP40* from BDSC, stock number 42747, and is described in[10].
11. *P{hsFLP}1* from BDSC stock number 6, and is described in[39]
12. *P{$\alpha$Tub84b(FRT.Gal80)}2* from BDSC, stock number 38880, and is described in[5].

**Fly Husbandry:**

All flies were raised on molasses-based food at 25°C with the exception of CsChrimson expressing flies, which were raised on Nutri-Fly German Food (#66-115, Genesee Scientific) food containing all-trans-retinal. Conditions for specific experiments are described below.

**Brain Dissection, Immunolabeling and Confocal imaging:**

Note: brain dissection, immunolabeling and confocal imaging were performed in two different laboratories with slightly different protocols.

**For Fig. 1, S3 and S4 (Clandinin Laboratory):**

Brain dissections were performed on 4-6 day old adult female flies. Flies were ordered in a fly collar, the proboscis and antennae were removed for each fly and flies were perfused with freshly-made 2% PFA in Phosphate-Buffered Lysine for 50min. We removed the fixative and washed brains 3X with ice-cold PBS and then proceeded to extract brains with fine forceps. Brains were stored in ice cold PBS +

0.1% Triton-X (PBS-Tx) for up to 2 hours and then were moved to a PBS-Tx + 10% Normal Goat Serum (NGS) blocking solution for 30min at room temperature. We incubated the brains in the following primary antibodies for 3 days at 4ºC: anti-GFP (chicken, Abcam, 1:2000), anti-Bruchpilot (nc82, mouse, Developmental Studies Hybridoma Bank, 1:30), anti-DsRed (rabbit, Clontech, 1:700). The anti-GFP antibody recognized both GFP and GCaMP6f. The anti-DsRed recognized myr-tdTomato. After primary incubation, brains were washed 3 x 15min in PBS + 0.1% Triton-X and then subjected to secondary staining for 2hrs at room temperature. All secondary antibodies were diluted 1:200 in PBS-Tx + NGS; these included anti-Chicken-alexa488 (Life Technologies, Carlsbad, Ca, USA), anti-rabbit-Cy3 (Life Technologies, Carlsbad, Ca, USA), and anti-mouse-Alexa-633 (Life Technologies, Carlsbad, Ca, USA). Brains were then washed 3 x 15min in PBS + 0.1% Triton-X, incubated for at least 1hr in 70% glycerol for tissue clearing, and mounted individually on glass slides in Vectashield (H-1000, Vector Laboratories) for confocal imaging.

Brains were imaged on a Leica SP8 confocal microscope. Series of between 20 and 100 optical sections (1.0 µm spacing) were imaged using either a Leica HC PL APO 20x/0.70 CORR CS oil-immersion lens (N.A. 1.3) or a Leica HC PL APO 40x/1.30 CS2 40x oil-immersion lens (N.A. 1.42). Tiffs of single confocal planes or maximum intensity projections (MIP) were made in Imaris 9.3 (Oxford Instruments, Abingdon, UK, GB). For Fig. 1 and Fig. S4I-K, MIP were from 3 sections (3um) of tissue, for Fig. S4A-D, MIP of Mushroom Bodies were generated from 15-20 5um optical sections of, while images of cell bodies (T4 and T5, Fig. S3; Kenyon cells, Fig. S4E-H) were taken from single optical sections.

**Image Processing**

Tiffs of single confocal planes or MIPs were generated in Imaris 9.3 and subsequently rotated and cropped to the same dimensions using Adobe Photoshop. Brightness levels were uniformly adjusted in Adobe Photoshop across images that were compared within a figure.

**Cell Counting**

T4 and T5 (Fig. 1 and S3) and Mi1 (Fig. S4) cell bodies were imaged at 40X using a series of 10 optical sections spaced 1$\mu$m apart as described above. Subsequently, single planes from the top, middle and bottom of these stacks were isolated for each optic lobe (N= 6 per condition) and individual tiffs were generated for myr-tdTomato and GCaMP6f stains from these planes. We randomly shuffled these images and a blinded author manually counted the individual cell bodies in each channel in Adobe Photoshop.  We calculated the percent of SPARC-labeled cells as total number of SPARC-labeled cells/total number of cells labeled in the Gal4 pattern by myr-tdTomato.

**For Fig. 2 (Wilson Laboratory):**

For immunostaining brain dissections, newly eclosed female flies that were raised on 0.6 mM all-trans-retinal-containing Nutri-Fly German Food (#66-115, Genesee

Scientific) were collected on $CO_2$. The brains were then dissected out of the head in chilled external saline[40]. Immunostaining was then performed as follows. Brains were (1) fixed in 4% paraformaldehyde (15714, Electron Microscopy Sciences) in phosphate buffered saline (PBS; 46-013-CM, Thermo Fisher Scientific) for 15 minutes at room temperature; (2) washed 3 times for 15 min with PBST (PBS with 0.44% Triton X-100 (T-8787, Sigma-Aldrich)); (3) incubated in a blocking solution of 5% normal goat serum (NGS; G9023, Sigma-Aldrich) in PBST for 20 min; (4) incubated in a primary antibody solution containing mouse anti-Bruchpilot antibody (1:25, nc82, Developmental Studies Hybridoma Bank), chicken anti-GFP (1:1000, ab13970, Abcam), and rabbit anti-dsRed (1:500, 632496, Clontech) diluted in the NGS blocking buffer for 48 hr at room temperature on a rotating nutator; (5) washed 3 times for 15 min with PBST; (6) incubated in a secondary antibody solution containing Alexa 488-conjugated goat anti-chicken (1:250, A-11039, Thermo Fisher Scientific), Alexa 568-conjugated goat anti-rabbit (1:250, A-11011, Thermo Fisher Scientific), and Alexa 633-conjugated goat anti-mouse (1:250, A-21050, Thermo Fisher Scientific) in blocking solution for 24 hr at room temperature on a rotating nutator; and (7) washed 3 times for 15 min with PBST.

Brains were mounted on slides in Vectashield (H-1000, Vector Laboratories) in the anterior-posterior orientation and then imaged using a Leica SPE confocal microscope. Series of between 30 and 100 optical sections (1.0 $\mu$m spacing) were imaged using either an Olympus UPLFLN 40x oil-immersion lens (N.A. 1.3) or an Olympus PLAPON 60x oil-immersion lens (N.A. 1.42). MIPs of the cell body images were made in Fiji[41], and MIPs of the full R2 neuron morphology were made in Imaris 9.3 (Oxford Instruments).

**Fly Preparation for 2-Photon Imaging:**

For two-photon imaging of T5, we used the enhancer fragment VT025965 to drive expression of Gal4 in T5 with high specificity[37]. Sparse expression of GCaMP6f was achieved using either a FlpOut or SPARC strategy. For the FlpOut strategy, we used the genotype *+/yw, P(hsFLP); P(20XUAS-IVS-GCaMP6f)attP40/ P(αtubP(FRT.stop)Gal80)2; P(VT025965-Gal4)attP2 /*+ with heat-shock at 37°C for 90 seconds during the late 3rd instar stage of development[16,42]. This heat shock protocol yielded the sparsest expression pattern for this T5 driver. Shorter or developmentally later heat shock resulted in either no GCaMP6f expression, or no observable GCaMP6f signals. For the SPARC strategy, we used the genotype +*; UAS-SPARC-S-GCaMP6f / P(nSyb-PhiC31)attP5; P(VT025965-Gal4)attP2/* +

All flies were female, and were imaged within 5-7 days of eclosion. To prepare for imaging, flies were immobilized by ice, and affixed to a custom-built mount with UV-cured optical epoxy (NOA 63, Norland Optical Adhesives). The cuticle, fat bodies, and trachea of the left hemisphere were removed under ice-cold, artificial hemolymph without calcium[40] to expose the brain for imaging from above. During imaging, standard, carbogen-gassed, room-temperature artificial hemolymph[40] was perfused across the brain at 150 mL/h.

**Imaging and Delivery of Visual Stimuli:**

Imaging and delivery of visual stimuli follow Leong et al. 2016[17]. Fluorescence was monitored *in vivo* using two-photon microscopy. For two-photon calcium imaging, we used a Leica SP 5 II equipped with a 20X 1.0NA lens (GCaMP6f excitation @ 920 nm, ~5-8 mW at the stage). Recordings lasted ~3.5 minutes. GCaMP6f fluorescence signals were acquired with a bandpass filter (525/50m), at ~20 Hz (bidirectional scanning at 1.4 kHz, across a FOV of 128 pixels x 256 pixels, rows x columns). Pixels measured ~290 x ~290 nm. The stimulus screen subtended ~ 60° x 90° (azimuth x elevation) of the left visual field. Visual stimuli were delivered with a Lightcrafter 4500 DLP, configured to deliver exclusively blue LED illumination, using a 100 Hz frame rate. The stimulus was attenuated with a 447/60 bandpass filter (Semrock), and a ND1 filter (Thorlabs). The mean radiance was ~0.04 W $sr^{-1}$ $m^{-2}$.

**Identification and Selection of ROIs:**

ROI selection involved two stages: (1) automated segmentation[43] of GCaMP6f responses to moving sinusoidal gratings to obtain an initial set of ROIs, each representing approximately individual cells (2) exclusion of ROIs from this initial set if they did not match the known calcium response properties of T5[16,17], or if their spatiotemporal receptive fields did not lie in the center of the stimulus screen, yielding a final set of ROIs that best represent individual T5 dendritic arbors. As T5 dendrites are fine and interdigitating, this ROI selection strategy could not always isolate dendritic arbors of individual T5 cells, particularly for FlpOut clones, since the sparsest possible FlpOut expression pattern that we could achieve was denser than the sparsest possible SPARC expression pattern. GCaMP6f responses to moving light and dark edges were used to confirm dark contrast selectivity (data not shown), and the timing of responses to moving dark edges was used to determine whether the cell's spatiotemporal receptive field was centered on the stimulus screen (data not shown). ROIs that did not meet these criteria were not imaged.

**Experiment, Stimulus Design, and Data Analysis:**

We presented sinusoidal gratings (1Hz, 25°/cycle, 100% contrast) moving for 5 seconds in 8 equally-spaced directions. Each 5-second bout was preceded by a 3-second "blank" (a gray screen, of luminance matching the mean luminance of the gratings). We presented 3 complete cycles of all 8 directions, in random order, for a total recording duration of ~3.5 minutes.

GCaMP6f fluorescence responses were quantified as the average $\Delta F/F_0$ across pixels within each ROI, where $F_0$ was defined as the mean fluorescence within each ROI during the final 5 frames of the "blank" preceding each bout. Responses were averaged across all three bouts to obtain the mean response to each direction of motion (plotted in Fig. 2C). Tuning curves (Fig. 2D) were derived from these mean responses, plotted as the maximum $\Delta F/F_0$ for each direction, normalized by the maximum $\Delta F/F_0$ across directions (the PD response), and registered across ROIs to align the PD response before averaging across ROIs. For each ROI, DSI (Fig. 2E) was calculated as the vector average of response amplitudes to the 8 directions of motion, normalized by the sum of response amplitudes to all 8 directions of motion.

**Fly preparation and dissection for electrophysiology:**

Newly eclosed virgin female flies were collected on ice approximately 1-4 hrs before the experiment. All flies were raised on 0.6 mM all-*trans*-retinal-containing Nutri-Fly German Food (#66-115, Genesee Scientific) and fly vials were wrapped in foil to prevent photo-conversion of the all-trans-retinal. At the beginning of each dissection, the fly was cold-anesthetized.

The preparation holder consisted of a flat titanium foil secured in an acrylic platform, with the foil oriented parallel to the horizontal body plane; the fly's head and body were gently pushed partway-through a hole in the foil. The head was pitched backward so that the anterior surface was oriented dorsally in the holder. The fly was always secured in the holder with epoxy (Loctite AA 3972) cured using a brief (<1s) pulse of UV light (LED-200, Electro-Lite Co). After the dorsal portion of the head was covered in saline, a large hole was cut in the head capsule and the retina and trachea on one side of the brain were removed to expose the neurons of interest. To reduce brain movement, muscle 16 was severed, and the proboscis was removed. An aperture was made in the perineural sheath around the somata of interest by ripping gently with fine forceps.

The external solution contained (in mM): 103 NaCl, 3 KCl, 5 N-tris(hydroxymethyl) methyl-2-aminoethane-sulfonic acid, 8 trehalose, 10 glucose, 26 $NaHCO_3$, 1 $NaH_2PO_4$, 1.5 $CaCl_2$ and 4 $MgCl_2$, with osmolarity adjusted to 270–273 mOsm. External solution was bubbled with 95% $O_2$ and 5% $CO_2$ and reached a final pH of 7.3. External solution was continuously perfused over the brain during electrophysiology.

**Whole-cell patch clamp recordings:**

*In vivo* whole-cell patch clamp recordings were performed as described previously[44]. Patch pipettes were made from borosilicate glass (1.5mm O.D., 0.86 I.D., # BF150-86-7.5HP, Sutter Instrument Co.) using a Sutter Instrument P-97 puller. Pipettes resistances ranged from 5-12 MΩ. The internal solution contained (in mM): 140 potassium aspartate, 10 4-(2-hydroxyethyl)-1-piperazineethanesulfonic acid, 4 MgATP, 0.5 $Na_3GTP$, 1 ethylene glycol tetraacetic acid, 1 KCl, and 13 biocytin hydrazide. The pH was 7.3, and the osmolarity was adjusted to ~268 mOsm. Recordings were performed at room temperature.

To obtain patch-clamp recordings under visual control, we used an Olympus BX51WI microscope with a 40X water-immersion objective (LUMPlan FI/IR NA 0.8, Olympus). GFP and tdTomato expressing neurons were identified using an Hg-lamp source (U-LH100HG, Olympus) with an EGFP-longpass filter (U-N41012, Chroma) or a TRITC-Cy3 filter (Chroma).

To visualize the brain for recordings, far-red light was delivered from a fiber-coupled LED (740nm, M740F2, Thorlabs) via a ferrule patch cable (200 µm core, Thorlabs) plugged into a fiber optic cannula (Ø1.25 mm SS ferrule 200 µm core, 0.22 NA, Thorlabs) glued to the recording platform, with the tip of the cannula ~1 cm behind the fly.

Recordings were obtained using an Axopatch 200B amplifier and a CV-203BU head-stage (Molecular Devices). Voltage signals were low-pass filtered at 5 kHz prior to digitization and then acquired with a NiDAQ PCI-6251 (National Instruments) at 20 kHz. Liquid junction potential correction was performed *post hoc* by subtracting 13 mV from recorded voltages[45]. When a stable whole-cell recording was achieved, the initial resting membrane potential was measured. Consistent with what one might expect from expression of a cation channel, we observed differences between the resting membrane potential and input resistance between cells expressing CsChrimson and control cells:

| R2 neuron transgene Expression | Resting Membrane Voltage (mean ± SEM, mV)* | Input Resistance (mean ± SEM, MΩ)* |
|---|---|---|
| CsChrimson::TdTomato+, GFP+ | -38 ± 5 | 220 ± 14 |
| GFP+ | -48 ± 2 | 628 ± 131 |

* P<0.05, Student's t-test

During optogenetic stimulation, a constant hyperpolarizing current was applied to bring the cell's membrane potential to between -50 mV and -60 mV.

For optogenetic stimulation, the Hg-lamp source (U-LH100HG) was used to deliver a 50-ms pulse of green light (530-550 nm, 2-4 mW, TRITC-Cy3 filter cube, Chroma) via the objective. A shutter (Uniblitz Electronic) controlled the pulse duration.

**Electrophysiology data analysis and data inclusion:**

To measure CsChrimson responses, the mean of 10 replicate stimulation trials was taken and filtered using a median filter with a 20-ms window to remove the effect of spiking activity. Evoked response amplitudes were the largest deviation from baseline that occurred within the 500 ms following the optogenetic stimulation. The one second preceding stimulation was used as the measurement of baseline membrane voltage.

Cells were only analyzed if the resting membrane voltage of the cell was < -30mV immediately following break in. One of eight recordings were excluded based on this criterion.

**Code availability**

All analysis was carried-out using custom-written MATLAB code. Visual stimuli were programmed with the OpenGL 1.0 API in Visual C#. All code is available on Github and will be made available upon request from the corresponding author.

**Data availability:**

All data will be made available upon request from the corresponding authors.
